# Non-invasive ultrasonic modulation of visual evoked response by GABA delivery through the blood brain barrier

**DOI:** 10.1101/351270

**Authors:** C. Constans, H. Ahnine, M. D. Santin, S. Lehericy, M. Tanter, P. Pouget, J-F Aubry

## Abstract

We demonstrate the feasibility of non-invasively modulating the visual cortex activity of non-human primates by local ultrasound-induced delivery of an inhibitory neurotransmitter (GABA). GABA was injected intravenously after the blood brain barrier (BBB) was transiently disrupted with focused ultrasound (FUS) coupled with ultrasound contrast agents (UCA). Visual evoked potentials exhibited a significant and progressive decrease of the activity. Combined effects of neuromodulation and BBB opening were shown to be 8.7 times less important than GABA-induced inhibition. During the sonication, the UCA harmonic response was monitored to estimate the level of stable cavitation (signature of BBB opening efficiency) and to avoid damages due to inertial cavitation (automatic shutdown of the sonication when detected). As recent developments in beam forming have shown that ultrasound beams can be focused non-invasively in deep-seated human brain locations, our results hold promise to explore and treat the brain with a non-invasive, controllable, repeatable and reversible method.

## Introduction

The blood brain barrier (BBB) naturally prevents large molecules (more than 0.4 to 0.5 kDa) from diffusing through the capillary walls of the brain^1,2^. While protecting the brain from numerous toxic agents, it also prevents potential medication for brain diseases to being administered through the blood^3^. The BBB can be temporary and reversibly lifted^4,5^ through the intravascular injection of microbulles coupled with low-intensity ultrasound^6,7^: the acoustic wave induces the bubbles oscillations in the fine brain capillaries, leading to the temporary disruption of the cohesion of endothelial cells through tight junctions which ensured the BBB efficiency. The BBB opening lasts a few hours^8^ and its safety has been investigated in several studies on small animals^9–13^ and non-human primates^14,15^, highlighting the possibility of non-destructive, efficient BBB openings with a mechanical index (MI) below 0.46 in rabbits^16^ and 0.58 in monkeys^14^. None of these early studies have shown functional consequences of BBB opening.

The technique holds promise for therapeutic drug delivery. A clinical trial has been conducted with the objective of delivering chemotherapy on patients with glioblastoma by opening the BBB with an ultrasound device implanted into the skull^17^. The technique could also provide a novel tool for non-invasive and local brain modulation by delivering inhibiting of stimulating drugs. Current neuromodulation techniques have improved dramatically in the last decades but present incompressible drawbacks: deep brain stimulation^18,19^ and localized injection of neuroactive substances are invasive^20^, optogenetic methods cannot be applied to humans, direct current stimulation^21^ and transcranial magnetic stimulation^22–24^ have a low spatial resolution^25^ and are limited to cortical brain areas^22,26^. Focused Ultrasound (FUS) techniques are rising as non-invasive, localized neuromodulation tools that can be potentially applied to humans. Modulation with ultrasound alone has been exhibited in both animals and humans. The observed effects are interestingly varied: modulation of EEG response^27,28^, excitation of neuronal circuits^29,30^, modulation^31^or elicitation^32^ of sensory sensations, behavioral changes^33,34^, motor response^35–39^, suppression of the somatosensory evoked potential (SSEP)^40^ and enhancement of neurogenesis^41,42^. However, as the mechanism of FUS-induced neuromodulation is still not fully understood^43,44^ the list of its effects is possibly not complete and as for today, the technique is more in a state of research than a tool for brain mapping or planned modulation. Nevertheless, the advantages of precise ultrasound focusing could be combined with the predictability of neuroactive agents’ action with BBB opening. For example γ-Aminobutyric acid (GABA) is a neuroactive agent which does not normally pass through the BBB^45^. McDannold et al. (2015)^46^ demonstrated the feasibility to temporary suppress the SSEP in rats by GABA delivery with ultrasonic BBB opening. Zhang et al. (2016)^47^later disconnected rats brain circuitry by introducing quinolinic acid after magnetic-resonance guided ultrasonic BBB opening. Here we demonstrate that this technique is non-invasive, controllable, repeatable and reversible on anesthetized non-human primates with a real-time monitoring of bubbles harmonic response to ensure both safety and efficiency of the BBB opening. For the first time, functional modulation induced by BBB opening is observed in non-human primates. We targeted the visual cortex of the animals with a single-element transducer operated at 245 kHz. We observed a decrease of the visual response intensity to full field visual stimuli and investigated the GABA dose dependency of this effect. The BBB opening was confirmed twice with MRI acquisition. Our goal was to evaluate the relative impact of FUS alone, FUS with UCA, and GABA delivery on the visual response.

## Results

### MRI

To verify the efficiency of the FUS system, we performed two BBB openings on an anesthetized animal before an MRI acquisition. During these experiments, no visual evoked potentials (VEPs) were recorded but the FUS + UCA procedure was identical. We used gadolinium (gadoterate meglumine, DOTAREM^®^, Guerbet, France) as the MR contrast agent (MRCA). Gadolinium has a molecular weight of 938 Da and does not normally pass the BBB^8^. The diffusion of the MRCA in the brain tissue indicated where the BBB was disrupted.

Both BBB openings proved successful on the images: the MRCA appeared in the occipital and cerebellar areas where ultrasound was focused, after the FUS procedure only. Figure 1 displays the MRI images before and after BBB opening. The signal intensity of turbo spin echo T1-weighted images varied from 1.23 to 2.27 in the targeted region, relatively to a Region of Interest (ROI) defined within neck muscle.

**Figure 1:**
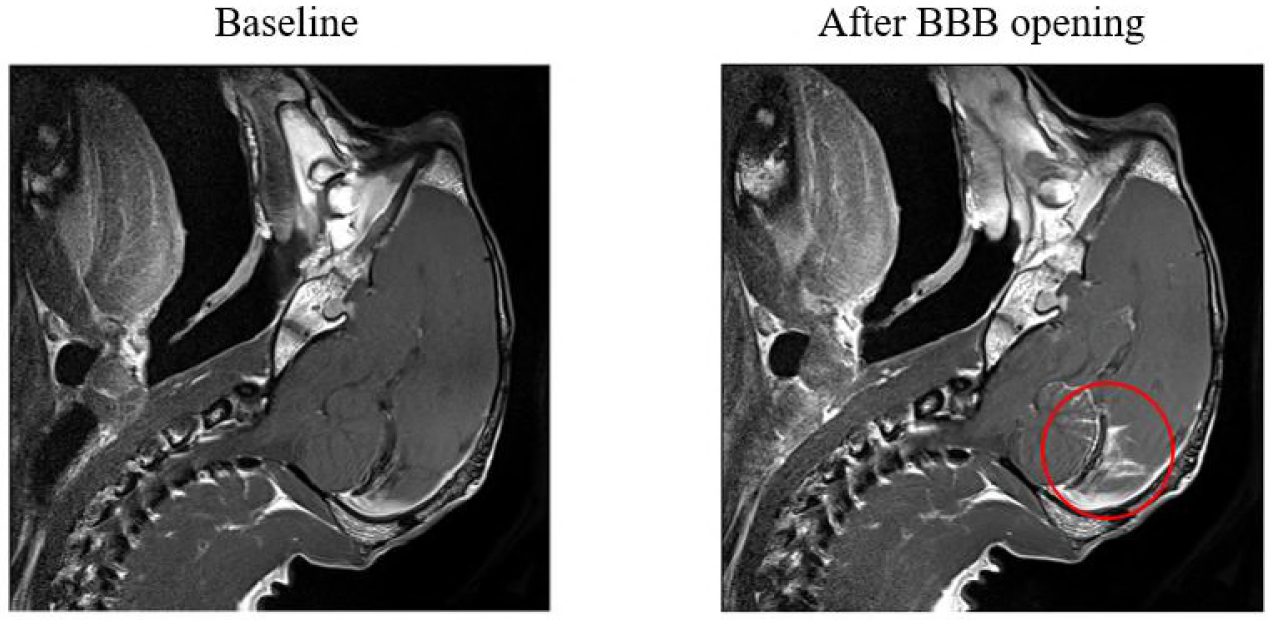
MRI assessment of BBB opening. Both images were obtained 10 minutes after injection of 2mL of a 0.5mmol /mL Gd. Left: before BBB opening. Right: after BBB opening. The red circle indicates the region where MRCA appeared.

### Response to visual stimuli

Figure 2 represents the timeline of the procedure. Three ‘’baseline’ runs were performed after installation of the animal and placement of the transducer, prior to any sonication. The animals were installed in sphinx position in front of a black screen. Eyes were kept opened. A run of visual stimuli consisted in 200 full field white flashes with a stimulus onset asynchrony (SOA) of 2s. The animals were placed in a dim room and we waited at least five minutes after the complete extinction of light in the room to start the first run. A ‘neuromodulation’ sonication was then launched, with the same ultrasound sequence than for BBB opening but without any injection of UCA. The ‘neuromodulation’ run occurred at the end of this sonication. UCA injection was then performed under the reduced light of a smartphone, via a catheter inserted in the small saphenous vein before light extinction. The ultrasound sequence coupled with the UCA injection was launched, followed by a ‘BBB opening, no GABA’ run. Finally GABA was injected intravenously (0.1 to 6 mg/kg) and at least 3 ‘GABA’ runs were conducted, depending on the animal temperature state.

Figure 2: Timeline of the experiments. Each bar represents a VEP run (measurement of the visual responses to 200 full field flashes)

Figure 3 displays the VEP results of one session (monkey A, GABA dose=5 mg/kg). Each curve represents the mean of 200 responses. The ‘baseline’ curve corresponds to the third baseline run, the first two runs being considered as a period of stabilization and dark adaptation (a total time of 20 minutes). The legend describes the runs in the chronological order. A FUS procedure without UCA is performed between the ‘baseline’ and ‘neuromodulation’ runs. Then the BBB is opened with FUS and UCA injection between the ‘neuromodulation’ and ‘No GABA’ runs. Finally, GABA is injected intravenously after the ‘no GABA’ run and the ‘GABA’ runs are conducted successively, each one lasting about 5 min. Sham sessions were performed without any ultrasound sonication nor UCA and GABA injection, but the timing of VEP recordings was identical to a non-sham session. The runs’ names were kept similar to the non-sham sessions (‘baseline’, ‘neuromodulation’, ‘no GABA’, ‘GABA’), even though there was no neuromodulation nor GABA injection.

**Figure 3:**
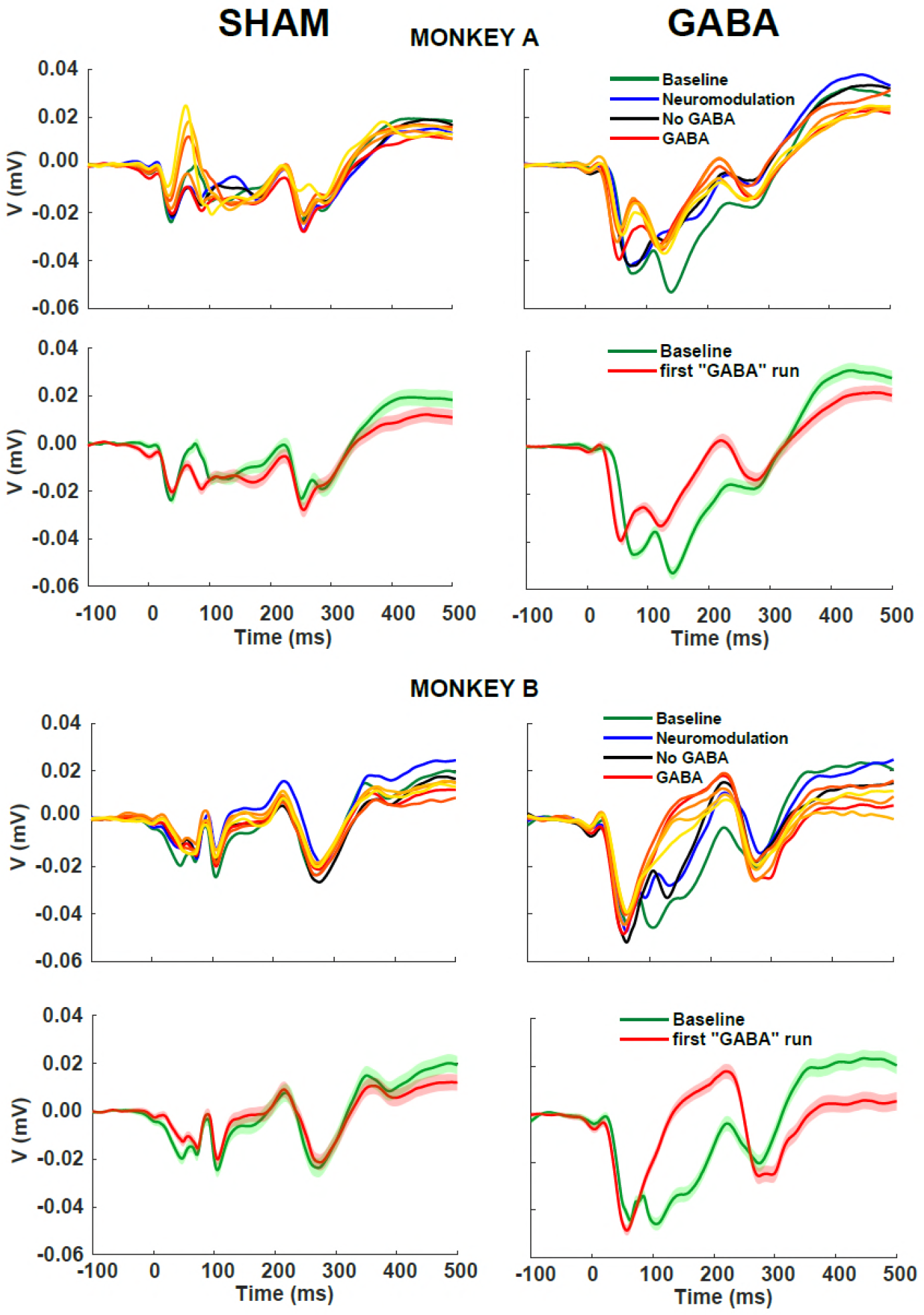
Mean VEP recordings for each run from baseline to ‘GABA 5’ for sham sessions on both monkeys (left), session 5 mg/kg GABA dose for monkey A and 4 mg/kg for monkey B (right). Each curve represents the VEP recordings of one run. The GABA runs are represented in orange gradient colors (red: GABA 1, yellow: GABA 5). The visual stimuli occur at time 0. For clarity purposes, each graph is replicated with only 2 runs (baseline run and first GABA run) with the standard error of the mean (SEM) (rows 2 and 4).

Another illustration of the decrease of visual response for both animals is shown in Figure 4. Only two of the recordings under sham conditions and at a 4 mg/kg GABA dose are displayed for each animal: the average of the VEP recordings of baseline 3 (before BBB opening), and the ‘GABA 1’ run (after BBB opening and GABA injection), corresponding to the first ‘GABA’ run.

**Figure 4:**
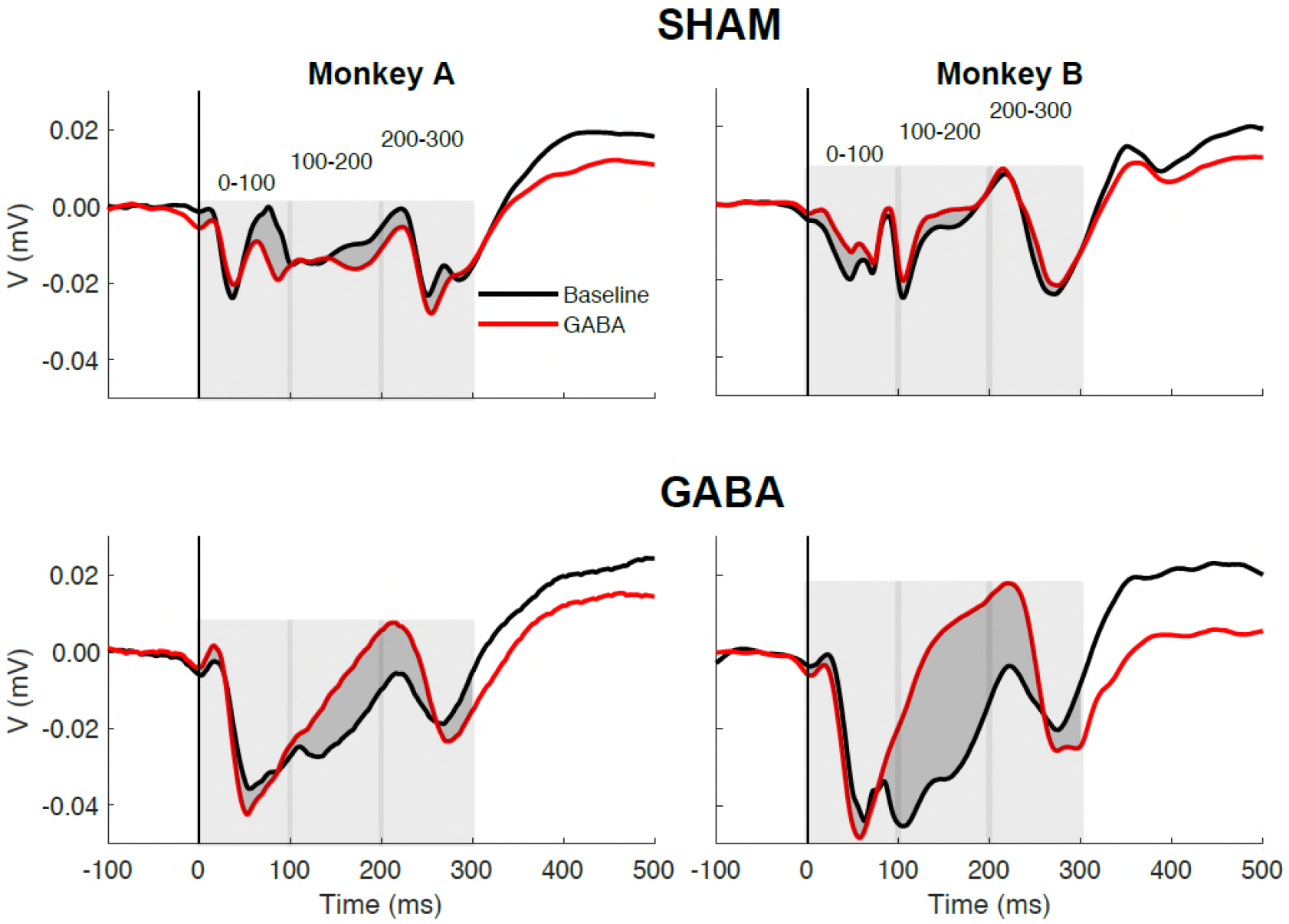
Comparison between the visual response before (Last baseline: black curves) and after (‘GABA 1’: red curves) neuromodulation by BBB opening and GABA injection for the two animals with a 4 mg/kg GABA dose (bottom) or under sham conditions (up). Each curve represents the mean of the 200 VEP recordings.

Finally, we performed a dose study. To quantify the decrease of the visual cortex activity during each session, we considered the decay of the VEPs P1 amplitudes by calculating the difference between the maximum and the minimum P1 peaks over all GABA runs. Figure 5 shows that the impact on P1 amplitude increases linearly when the GABA dose increases.

**Figure 5:**
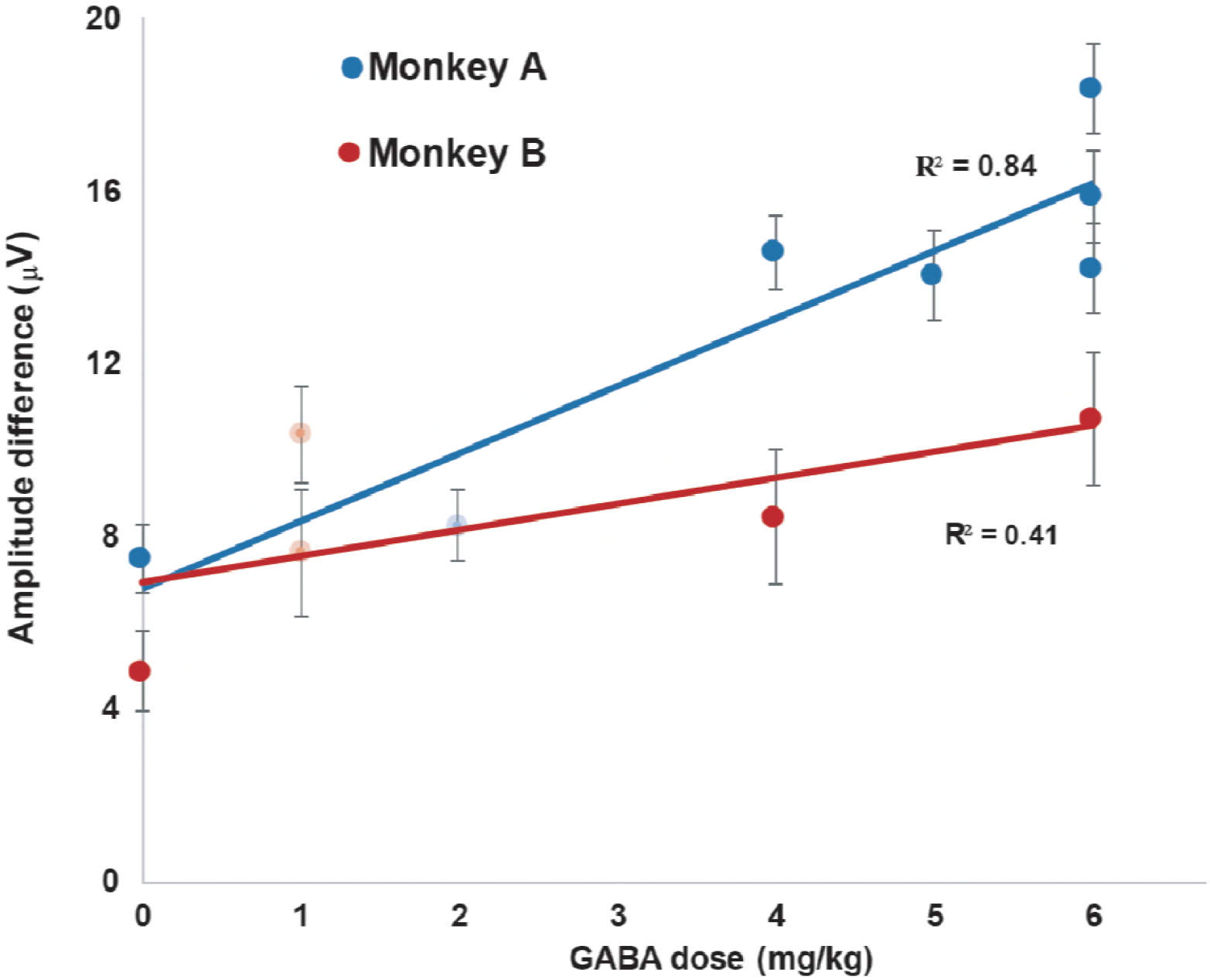
VEPs decrease of amplitude (maximum P1 amplitude – minimum P1 amplitude over the five first GABA runs (except for monkey A - 2mg and monkey B - 1mg with only 3 GABA runs, in light colors)) after GABA injection, as a function of GABA dose.

### UCA harmonics content

For every pulse, the signal received by the hydrophone, monitored on the second channel of the TiePie oscilloscope, was analyzed in the frequency domain. The level of broadband, harmonics, subharmonics and ultraharmonics was displayed in real time. The subharmonic is the signal at half excitation frequency f_0_/2 (here, 245/2 = 122.5 kHz). The subharmonic emission is known to be associated with stable cavitation^48,49^. Harmonics (f= n*f_0_ with n ≥2) and ultraharmonics (f= (n+1/2)*f_0_ with n ≥1) emissions, which are also often associated with stable cavitation^49^, were also recorded. The broadband emission corresponds to all the other frequencies. This noise, caused by the collapse of the bubbles, is related to inertial cavitation^50,51^.

The bolus injection of 2mL of UCA (SonoVue, Bracco, Milan, Italy) took place after the beginning of the sonications, typically between the 20^th^ and 50^th^ seconds (out of 200). A second injection of physiological serum with the same syringe was administrated a few seconds later to flush the rest of UCA that could have deposited in the syringe.

The levels of the different harmonics types and the broadband were first recorded prior to UCA injection. The relative augmentation 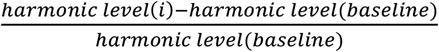 was calculated for every ultrasonic emission *i* for all types of harmonics and the broadband. An arbitrary safety threshold of 3 was set for the broadband maximum relative augmentation: if this value was reached, the sonication would be halted automatically. An efficiency threshold was also set to 3 for the harmonics and the subharmonic (f_0_/2) minimum relative augmentation as an indicator of BBB opening.

Figure 6 displays the relative change of harmonic level for each pulse (one per second) for the session with monkey A at 5mg/kg, which VEP results were presented previously (figure 2). The bolus injection of UCA started at shot #15 and ended at shot #29. The syringe was rinsed with a bolus of physiological serum between shots #47 and 50.

**Figure 6:**
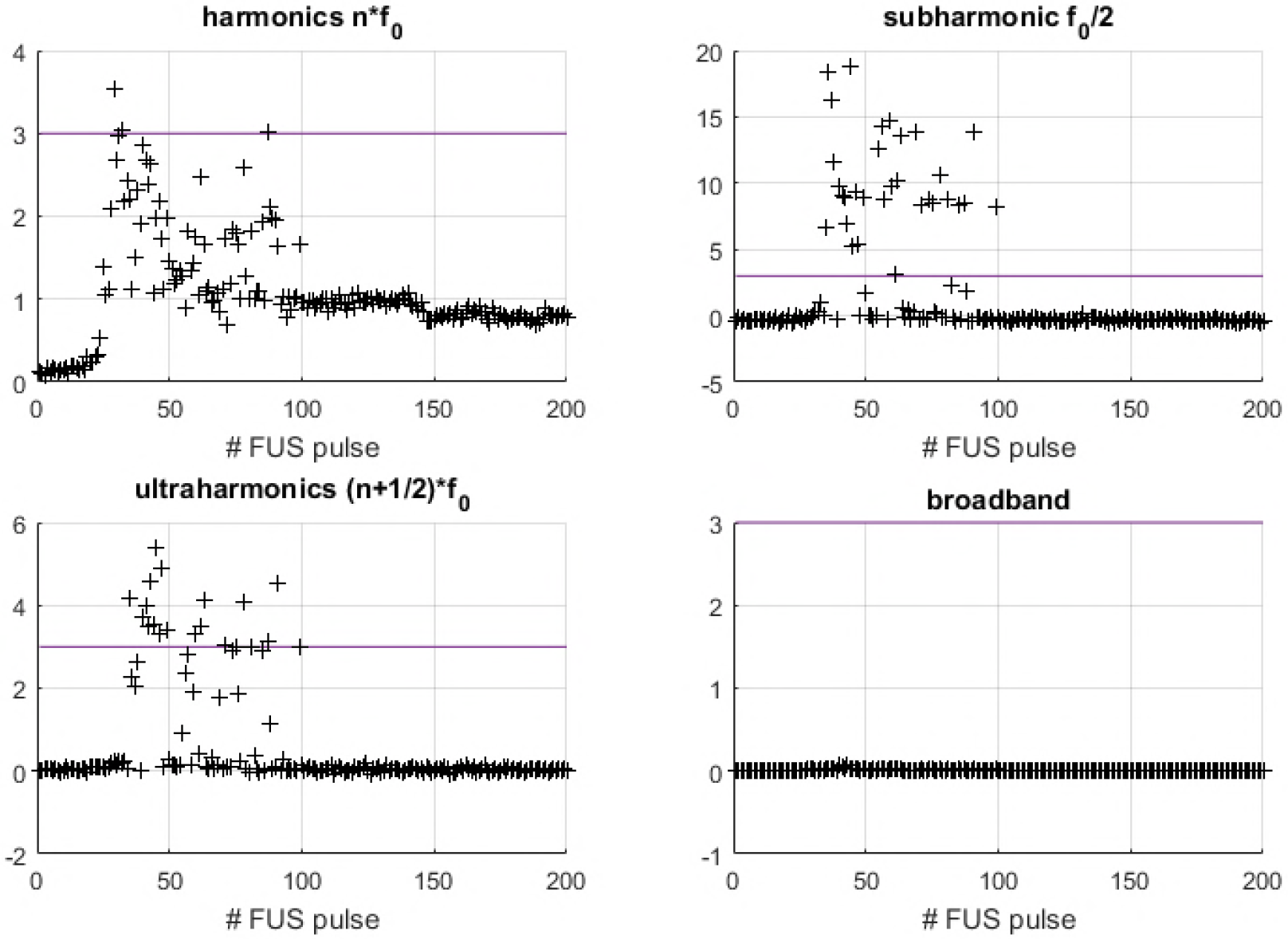
Harmonics and broadband level during BBB opening, session with monkey A at 5 mg/kg.

Figure S1 (supplementary materials) displays the mean harmonic content of the same session before the microbubbles injection (between the 1^st^ and 15^th^ pulses) and after (between the 45^th^ and the 60^th^ pulses). All types of harmonics are clearly emitted during the second period.

The subharmonic emission was above the efficiency threshold for at least 21 seconds for all sessions.

Table 1 (supplementary materials) shows, for each session, the maximum level of relative augmentation for each type of harmonics and broadband. We also calculated the time spent by the subharmonic level above the efficiency threshold.

## Discussion

Analysis of VEPs recordings showed a decrease of the visual response to full field flashes following the GABA dose (figures 3 and 5). The activity was not suppressed entirely, whereas McDannold et al. (2015)^46^ could almost completely inhibit the cortical primary somato-sensory (S1) activity in rats. Several hypotheses could explain this difference.

First, we probably did not reach the GABA dose required to inhibit completely the structures were the BBB was open. Since there is no pharmacology data on BBB disruption-induced GABA in the brain, the dose was limited to 6 mg/kg in our study for the safety of the animals (due to possible peripheral effect of GABA). For sake of comparison, McDannold et al. (2015)^46^ injected up to 60mg/kg in rats.

Second, even at a frequency as low as 245 kHz, the focal spot did not cover the entire area involved in the visual circuit, as it can be seen on the MRI (figure 1). In the rat study, BBB disruption was produced in both the cortex and subcortical structures such as the thalamus^46^, hence the possibility of a complete inhibition.

Third, the inhibition of the visual cortex might not be as straightforward as the inhibition of S1 cortex. The contributions of distinct primary visual areas to feedforward and feedback connections to the electrical potential recorded by VEPs are complex to disentangle. Based on peak latencies, an incremental delay between V1, V2, V3, and V3A visual latency has been reported, suggesting serial stages of processing. The extent to which early visual areas have distinct time courses of activation is, however, somewhat contentious^52,53^. According to direct recordings in monkeys, early visual areas first become active nearly simultaneously^54,55^. Additionally, V2, V3, and V3A receive some degree of direct, subcortical input that bypasses V1^56–61^.

Historically, single-cell recordings in nonhuman primates have shown that inactivation of higher-order areas modulates neuronal responses in lower-order areas^62–65^. It has been shown that V1 activity is modulated by GABA inhibition of area V2. Another study found similar results for V1, V2, and V3 neurons when area MT was inactivated^66^. Many studies indicate feedback signals mediating surround suppression of V1 neurons. Taken together these results strongly support the role of feedback from higher visual areas in determining V1 neural activity. Feedback interactions in human vision were also reported recently. It has also have been shown that early (40–100 ms) inactivation of V1, using transcranial magnetic stimulation (TMS), inhibits detection of simple features, but not conjunctions^56^. Conversely, inactivation of V1 after longer delays (200–240 ms) seems to impair detection of feature conjunctions, while leaving simple feature detection intact. This double-dissociation implicates V1 in feedback loops with higher visual areas, although it does not specify from where such feedback might originate. Other TMS studies^67,68^ specifically indicated feedback inputs from MT to V1 with latencies 80–125 ms from the stimulus onset.

Using transcranial ultrasound method, McDannold et al. (2015)^46^ raised the question of possible neuromodulation effects on the somatosensorial cortex activity during BBB opening, independently of the GABA inhibition. Indeed, several studies reported that ultrasound-induced neuromodulation can modulate brain function in primates^31,33,34,69^. In 5 out of 7 case, we observed a significant neuromodulation effects, before the GABA injection, between 100 and 300ms after the stimulus onset (figure 3 is one example). We therefore calculated the spectral power of the signals at each step: the baseline run, after ultrasound alone (‘Neuromodulation’ run), after ultrasound coupled with UCA before GABA injection (‘No GABA’ run), and finally after GABA injection (‘GABA’ run). We calculated the mean contribution of each step in the inhibition (corresponding to the spectral power decrease) over all sessions with a GABA dose of at least 4mg/kg for three different time periods: 0-100ms, 100-200ms and 200-300ms after the stimulus onset (figure 7). The neuromodulation contribution corresponds to the decrease of activity after the ‘neuromodulation’ run, i.e. the neuromodulation spectral power minus the baseline spectral power; the UCA contribution is the one after the ‘no GABA’ run, i.e. the ‘No GABA’ spectral power minus the neuromodulation spectral power; the GABA contribution is the one from the most perturbed ‘GABA’ run. In this analysis, we considered the spectral power instead of the amplitude in order to quantify the cerebral activity on different time periods after the stimulus onset. Results showed that FUS+UCA influence got stronger as time increases after the visual stimulus onset, compared to GABA-induced effects. The percentage of GABA-induced inhibition relative to the total inhibition (combined effects of neuromodulation, UCA and GABA) is 90% during the first 100 ms, 42% during the 100-200ms period and 50% during the 200-300ms period.

The translation of this work to the human anatomy requires further developments but will be facilitated by the recent development of multi-element transcranial ultrasound devices^70–74^, as well a low cost approach taking advantage of an acoustic lens to compensate for the aberrations induced by the human skull ^75^.

**Figure 7:**
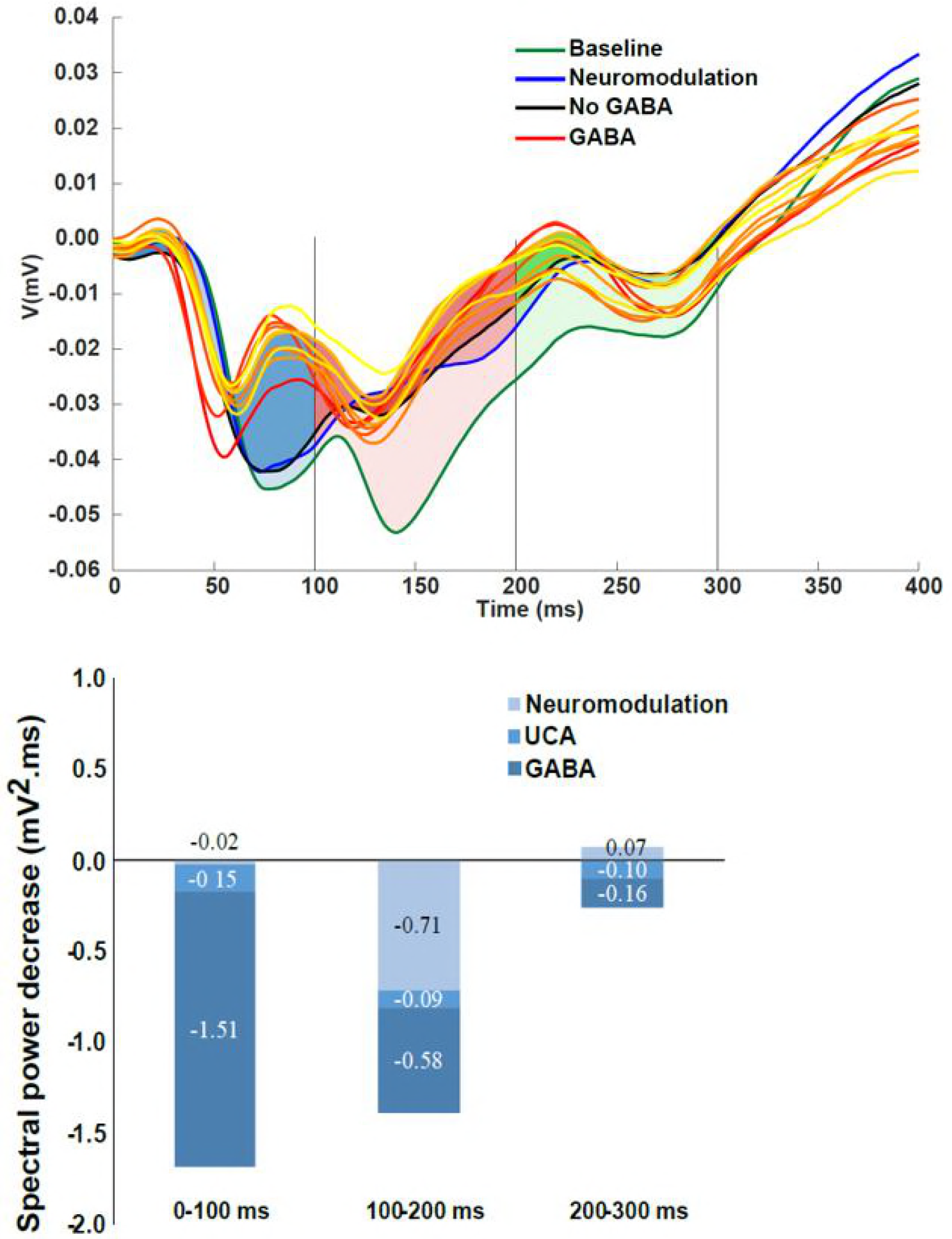
Proportion of the inhibitory effects due to neuromodulation (ultrasound only), BBB opening (ultrasound + UCA) and GABA (after BBB opening and GABA injection). Top: Illustration of the calculation of the contributions from the three events for one session (monkey A, GABA: 5 mg/kg). Bottom: Quantification (average of all sessions with a GABA dose of at least 4mg/kg on monkey A) of the contributions from neuromodulation, UCA and GABA on VEP spectral power decrease, for three different time periods.

## Materials and methods

### Animals

Two captive-born macaques (Macaca mulatta ‘A’ and ‘B’), both 6-year-old and respectively 8 and 10 kg, participated to the study. Monkeys were paired-housed and handled in strict accordance with the recommendations of the Weatherall Report about good animal practice. Monkey housing conditions, surgical procedures and experimental protocols were all carried out in strict accordance with the authorization for conducting experiments in our institute delivered by the Animal Health and Veterinary Medication Division of the Department of Public Veterinary Health, Nutrition and Food Safety of the French Ministry of Health (last renewal: Arrêté prefectoral N° DTPP B-75-13-19). Monkeys were enrolled in the project ‘Thérapie non invasive du cerveau par ultrasons focalisés’ (Non-invasive brain therapy using focused ultrasound) validated by the ethical committee C.DARWIN under the reference #6355. Our routine laboratory procedures included an environmental enrichment program where monkeys had access to toys, mirrors and swings. Monkeys also had visual, auditory and olfactory contacts with other animals and, when appropriate, could touch/groom each other. An institutional veterinary doctor regularly monitored the well-being and health conditions of the monkeys.

Anesthesia was induced with a blend of ketamine hydrochloride (3 mg/kg i.m.) and dexmedetomidine (0.015 mg/kg i.m.) for initial sedation and animals were anesthetized with isoflurane during the entire procedure (1.5% during installation, 1% during experiments). All procedures lasted less than 3 hours. Heart rate, temperature and respiration were monitored and kept within physiological range. The animal bodies were covered with survival sheets to limit the temperature decrease.

### Focused Ultrasound and Harmonics control

A single element FUS transducer (H117, Sonic Concept, Bothell, WA, USA) (center frequency 261kHz, diameter 64mm with 20mm central opening, F=1) with a passive cavitation detector (PCD) in its center (Y107, Sonic Concept, Bothell, WA, USA, 17.5mm active diameter, 64mm geometric focus, 10kHz to 20MHz bandwidth) was used at the frequency of 245 kHz. A coupling cone (C101, Sonic Concepts, Bothell, WA, USA) filled with degassed water was placed between the transducer and the animal head. The transducer was fixed on a mechanical arm with 4 rotation axes (Viewmaster LCD, Osmond Ergonomics, Wimborne, UK) to provide the flexibility for the positioning and orientation of the transducer over the head. The transducer was placed manually, targeting the middle of visual cortex V1, the tip of the cone being place as parallel as possible to the skull surface. A thin layer of echographic gel (Aquasonic 100, Parker Laboratories Inc., Fairfield, NJ, USA)) was applied on the shaved skin and on the membrane of the coupling cone to ensure acoustic coupling.

The signal (20ms pulse every second for 200 seconds) was created by a function generator (33250A, Agilent, Santa Clara, CA). A 75-Watts amplifier (75A250A, Amplifier Research, Souderton, PA) was then used to deliver the required power to the transducer through a matching network and the input voltage of the transducer was monitored using a voltage probe (P6139A, Tektronix, Melrose, MA) connected to an oscilloscope (Handyscope HS5, Tiepie Engineering, Sneek, The Netherlands). The amplifier gain was set to deliver an output voltage Vout=200V peak-to-peak to the transducer.

A calibration was conducted before the UCA injection: at a given amplifier gain, the amplitude of the signal generated by the first function generator was ramped up to 0.6V (with 0.02V steps), corresponding to approximately 215 V after amplification. The generator amplitude corresponding to the closest amplified voltage below 200V was chosen for the experiments. Different harmonics type responses were analyzed from the PCD recording.

The levels of the different harmonics types (harmonics n*f_0_, subharmonic f_0_/2 and ultraharmonics (n+1/2)* f_0_) and the broadband were recorded for this given voltage and used as baseline.

In order to estimate the peak pressure in the brain, a clean and degassed primate skull specimen (Maccaca Mulatta skull) was put in front of the transducer in a degassed water tank and the pressure at the focus was estimated using a heterodyne interferometer^76^. A heterodyne interferometer uses a laser beam to detect the vibration of a Mylar membrane induced by the ultrasound wave with. The amplitude of the vibration is then converted to pressure with high sensibility and a flat frequency response^77^.

The transmission of ultrasound through the degassed primate skull was assessed at 6 different points randomly chosen on the skull. The transmission was found to be 82% ± 6%. The in situ pressure delivered to the monkey brain transcranially was subsequently estimated at 0.54 ± 0.03 MPa.

The equivalent Mechanical index (MI) value is 1.1 with an Intensity Spatial Peak Pulse Average (ISPPA) of 9.7 W/cm² in the brain. By taking into account the pulse duration and pulse repetition frequency (respectively 20ms and 1Hz, corresponding to a 2% duty cycle) during the sequence, the Intensity Spatial Peak Time Average (ISPTA) is estimated to be 194mW/cm² behind the primate skull.

### Visual stimuli and VEP recordings

The animals were installed in a sphinx position in front of a black screen. Eyes were kept opened and gel (Ocry-gel, TVM, France) was applied to avoid eyes drying. A run of visual stimuli consisted in 200 full field flashes separated by 2s intervals. Two electrodes were inserted symmetrically in the skin above the V1 regions. The reference electrode was subcutaneously inserted in the eye brows and the ground electrode in the maxilla. The VEPs were recorded on a MAP system (Plexon Inc., TX, USA).

Sham sessions were performed without any ultrasound sonication nor UCA and GABA injection, but the timing of VEP recordings was identical to a non-sham session. The transducer was positioned on the animal head in order to reproduce the non-sham conditions but was turned off.

### MRI

We ran two experiments of BBB opening without the visual stimuli but with MRI assessment of Gadolinium (Gd) diffusion on monkey A. The MR contrast agent (MRCA) was gadoterate meglumine (Dotarem^®^, Guerbet, France). We used a dose of 2mL of a 0.5mmol/mL Gd solution. The animal head was maintained with a stereotaxic frame during the image acquisition. Three MR acquisitions were obtained during the experiment. The first MRI acquisition was performed at baseline before sonication without MRCA. The second MRI acquisition was performed before sonication and 5 to 10 minutes after injection of the MRCA (2mL), and the third acquisition was performed 15 minutes after sonication with a second MRCA injection 10 minutes after sonication. The second and third acquisitions were separated by a delay of about 25 minutes.

MRI was performed with a 3T magnet (Prisma, Siemens, Germany) using an 8-channel receive only head coil specifically designed for non-human primate experiments (Life Services LLC, USA). Images were acquired with a 2D-sagittal T1-weighted turbo spin-echo sequence with the following parameters: TR/TE: 689/11 ms, Echo Train Length: 4; voxel size: 0.4*0.4*1.5 mm^3^, averages: 8, 10 slices, acquisition time: 7min 20s).

### Data analysis

Electrophysiological data were post-processed using Matlab (MathWorks). The ground electrode signal was subtracted from the VEP recordings. The resulting signals were then filtered with a Savitzky-Golay filter (order 1, 21 ms frame length) and averaged (200 VEPs for a run). The offset, calculated as the mean of the first 50ms, was removed from each curve.

For the spectral power calculation, the signal was reduced to its period of interest (e.g. 0-100ms after stimulus onset) before the Fourier transform. The spectral vector was then squared and summed over its length to get the spectral power.

To standardize the analysis among the sessions, we always considered the five first GABA runs, even though we could acquire more data in some sessions. The exception is for Monkey A – 2mg/kg and Monkey B – 1 mg/kg sessions, in which we have only 3 ‘GABA’ runs.

In Figure 5, the error bars represent the root mean square of the corresponding runs SEM (the ones giving minimum and maximum amplitudes).

In figure S1, the average harmonic response over 15 ultrasound pulses is represented on a logarithmic scale as the Fourier transform of the time signals before and after UCA injection. The plots are normalized by the resonant response at f_0_ after UCA injection.

## Supplementary material

**Table 1.** Maximum relative augmentation and time above the efficiency threshold for the subharmonic.

| Monkey #<br>experiment<br>number | GABA<br>dose<br>(mg/kg) | Maximum relative augmentation |  |  |  |  |  |  | Time above<br>the<br>efficiency<br>threshold |
| --- | --- | --- | --- | --- | --- | --- | --- | --- | --- |
| | | broadband | $n*f_0$ | $f_0/2$ | $(n+1/2)*f_0$ | $f_0/3$ | $f_0/4$ | $f_0/6$ | |
| Monkey B | 1 | 1.2 | 0.6 | 2.3 | 1.2 | 5.3 | 1.9 | 2.0 | 21 |
| Monkey B | 1 | 0.83 | 7.5 | 40 | 16 | 13 | 8.6 | 3.5 | 70 |
| Monkey A | 2 | 0.57 | 6.0 | 25 | 16 | 12 | 7.9 | 2.1 | 140 |
| Monkey A | 4 | 0.08 | 1.5 | 46 | 3.2 | 0.7 | 0.4 | 0.3 | 150 |
| Monkey B | 4 | 1.2 | 4.5 | 4.0 | 5.7 | 5.7 | 3.5 | 2.5 | >118 |
| Monkey A | 5 | 0.07 | 3.5 | 19 | 5.4 | 0.1 | 0.2 | 0.2 | 65 |
| Monkey A | 6 | 0.23 | 3.5 | 5.3 | 14.6 | 4.9 | 6 | 1.3 | 56 |
| Monkey A | 6 | 0.2 | 4.9 | 38 | 8.7 | 0.7 | 0.9 | 0.3 | 27 |
| Monkey A | 6 | 0.8 | 12 | 59 | 23 | 16 | 11 | 6 | >157 |
| Monkey B | 6 | 1.1 | 8.1 | 80 | 21.1 | 7.1 | 5.7 | 3.1 | >135 |

**Figure S1:**
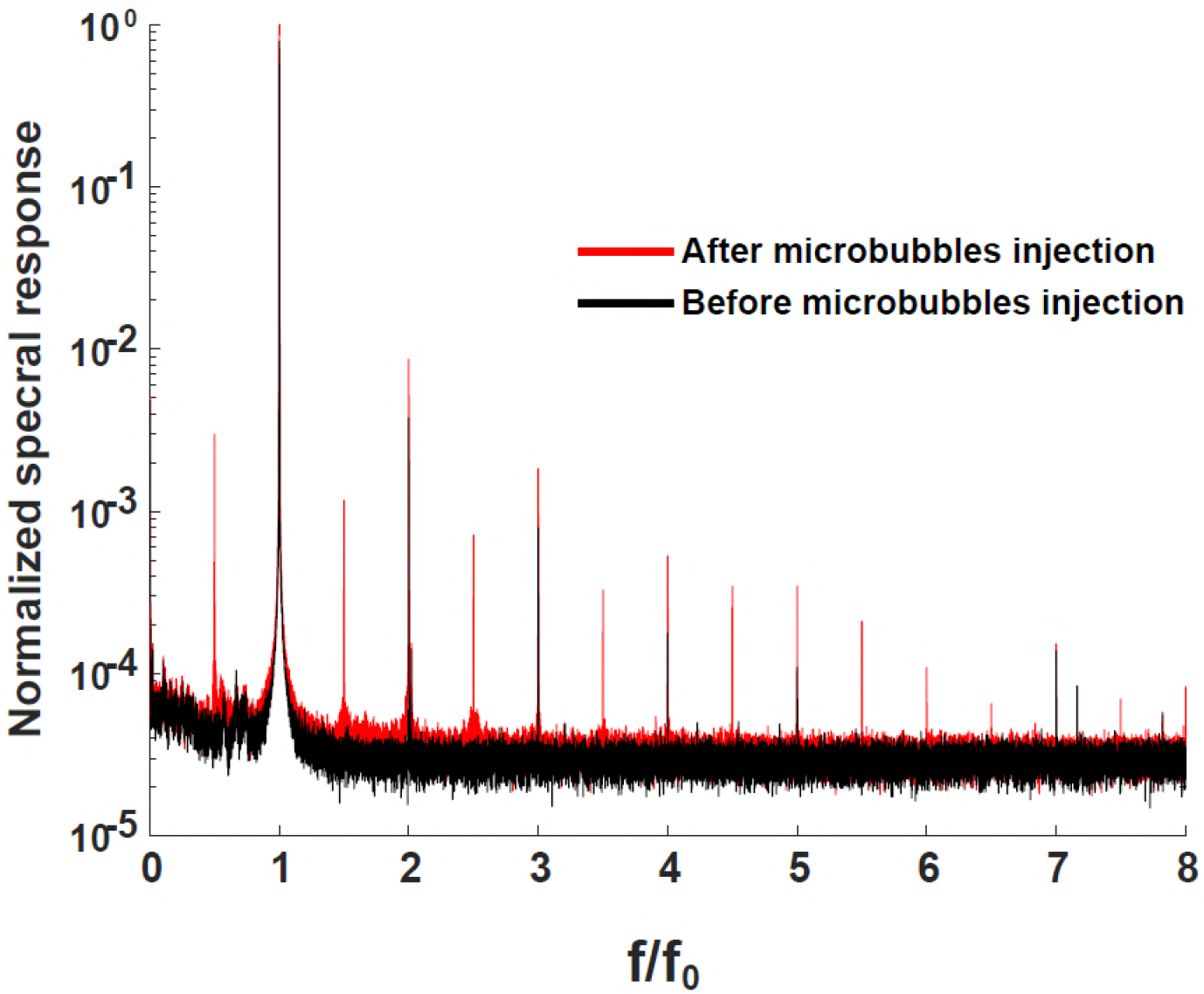
Spectral UCA response summed over t=1 to t=15 (before microbubbles injection) and over t=45 to t=60 (after microbubbles injection) on a logarithmic scale, session Monkey A 5mg/kg. The excitation frequency was f_0_=45 kHz.

## Acknowledgements

We thank Sophie Rivaud-Péchoux for her help on the statistical analysis.

This work has been supported by the Bettencourt Schueller Foundation, the LABEX WIFI(Laboratory of Excellence within the French Program ‘Investments for the Future’) under references ANR-10-LABX-24 and ANR-10-IDEX-0001-02 PSL, by the National Agency for Research under the program ‘future investments’ with the reference ANR-10-EQPX-15, IHU-A-ICM, Paris Institute of Translational neuroscience (IAIHU-06), France Life Imaging (ANR-11-INBS-0006), and NeurATRIS (ANR-11-INBS-0011 translational research infrastructure for biotherapies in neuroscience).

